# Classifying disease-associated variants using measures of protein activity and stability

**DOI:** 10.1101/688234

**Authors:** Michael Maglegaard Jepsen, Douglas M. Fowler, Rasmus Hartmann-Petersen, Amelie Stein, Kresten Lindorff-Larsen

## Abstract

Decreased cost of human exome and genome sequencing provides new opportunities for diagnosing genetic disorders, but we need better and more robust methods for interpreting sequencing results including determining whether and by which mechanism a specific missense variants may be pathogenic. Using the protein PTEN (phosphatase and tensin homolog) as an example, we show how recent developments in both experiments and computational modelling can be used to determine whether a missense variant is likely to be pathogenic. One approach relies on multiplexed experiments that enable determination of the effect of all possible individual missense variants in a cellular assay. Another approach is to use computational methods to predict variant effects. We compare two different multiplexed experiments and two computational methods to classify variant effects in PTEN. We distinguish between methods that focus on effects on protein stability and protein-specific methods that are more directly related to enzyme activity. Our results on PTEN suggest that ~60% of pathogenic variants cause loss of function because they destabilise the folded protein which is subsequently degraded. Methods that quantify a broader range of effects on PTEN activity perform better at predicting variant effects. Either experimental method performs better than the corresponding computational predictions, so that e.g. experiments that probe cellular abundance perform better at identifying pathogenic variants than predictions of thermodynamic stability. Our results suggest that loss of stability of PTEN is a key driver for disease, and we hypothesize that experiments and prediction methods that probe protein stability can be used to find variants with similar mechanisms in other genes.

## Introduction

Technological advances have made human genome sequencing feasible in routine clinical contexts, revealing around 10,000 changes in the protein-coding regions of each individual^1^, of which the majority will be very rare^2^ and difficult to interpret^3,4^. We use state-of-the-art high-throughput methods for assessment of protein variants and discuss their performance in discriminating pathogenic from non-detrimental changes. This preprint is a summary of a submitted manuscript.

PTEN (phosphatase and tensin homolog) is a two-domain protein (Fig. 1A and 1B) with dual-specificity phosphatase that acts on both protein and PIP_3_ lipid substrates. Its lipid phosphatase function is important for PTEN’s role as a tumor suppressor where it counteracts phosphatidylinositol 3-kinase function^5^. A number of germline mutations in the *PTEN* gene are associated with various diseases including autism spectrum disorder (ASD) and different tumor-risk syndromes collectively known as PTEN hamartoma tumor syndrome (PHTS)^5,6^ (Fig. 1B).

**Figure 1:**
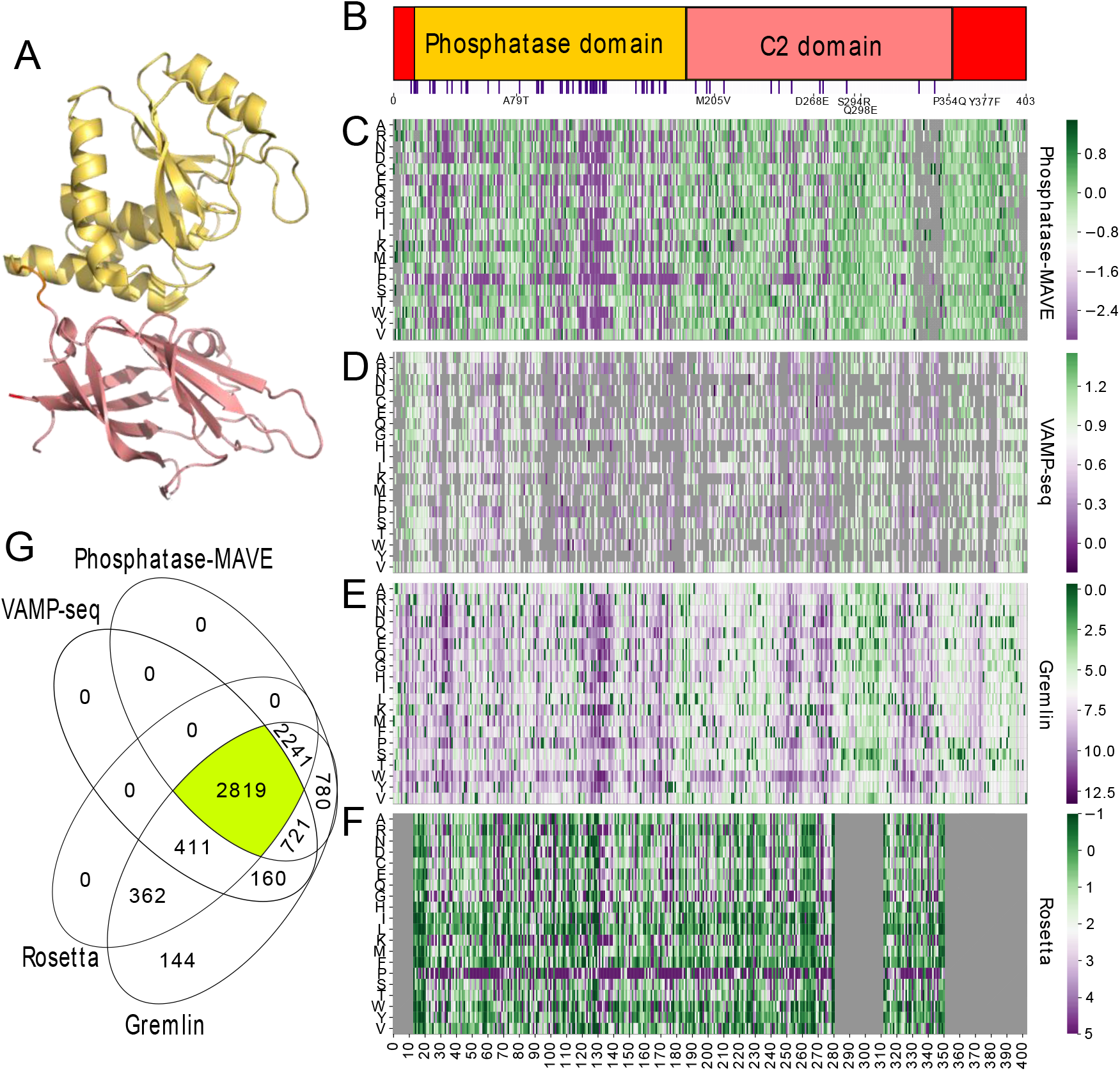
Overview of PTEN and the available single-variant data. A: We show the 3-dimensional structure of PTEN (PDB:1D5R (ref. ^7^)) with the phosphatase domain in yellow and the C2 domain in pink. B: PTEN domain architecture (top) and labels for gnomAD variants that are not singletons (bottom, details see main text). Purple bars indicate positions where pathogenic variants have been described. C-F: Experimentally and computationally determined scores for single site variants. Green indicates wild-type-like fitness/stability, magenta indicates low fitness/stability, gray indicates missing data. C: Phosphatase-MAVE^8^, D: VAMP-seq^9^, E: evolutionary sequence energies^10^, F: Rosetta cartesian ΔΔG (ref. ^11^). G: Venn diagram illustrating the availability of scores from each method and their intersections.

High-throughput genome sequencing combined with assays specific to the function of the protein of interest enable highly parallel assessments of the functionality of each individual variant^7^ in so called MAVEs (multiplexed assays of variant effects)^8^. Briefly explained, a MAVE involves creating a large library of variants and subsequently selecting for a property of interest^9–12^. MAVEs have been applied to a substantial number of proteins and domains, overall indicating a surprising mutational tolerance as many missense variants retain wild type-like function^13,14^.

In the case of PTEN, two different MAVEs have been performed using different selection systems. Mighell *et al*. evaluated the effects of PTEN variants on lipid phosphatase activity in a yeast system^15^ (Fig. 1C) and were able to discriminate likely pathogenic from benign alleles^15^. We term this experiment Phosphatase-MAVE. We and others have shown that loss of protein stability and subsequent degradation by the cellular protein quality control machinery is an important factor that underlie pathogenicity of variants in diverse proteins^16–21^. Matreyek *et al.* recently introduced VAMP-seq (variant abundance by massively parallel sequencing), as a MAVE of variant abundance^22^. Application of VAMP-seq to *PTEN* showed that many pathogenic variants were of low cellular abundance and that the resulting data (Fig. 1D) showed separation between pathogenic variants and common variants in the human population.

In the absence of experimental data, computational methods can be used to predict the consequences of missense variants. We here complement and contrast the experimental Phosphatase-MAVE and VAMP-seq experiments with two computational methods, one that assesses sequence conservation through the calculation of an ‘evolutionary sequence energy’ 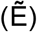 and one that quantifies changes in protein stability (ΔΔG). These methods have previously been applied to the identification and classification of pathogenic variants^4,23,24^. Loosely defined, 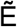-values are conceptually similar to the outcome of the Phosphatase-MAVE experiment and ΔΔG-values are related to the outcome of the VAMP-seq experiment. We use the software Gremlin^25^ to calculate evolutionary sequence energies (Fig. 1E) and the Rosetta software to calculate changes in protein stability^26^ (ΔΔG-values) (Fig. 1F). Out of the 7,638 possible single amino acids, 2,819 could be assessed by all four methods (Fig. 1G), however below we focus only on the smaller subset of 71 variants that are either known to be pathogenic (42) or that are found in the gnomAD database^27^ (29).

## Results and Discussion

As a source for pathogenic PTEN variants, we extracted 87 pathogenic missense variants from ClinVar^28^ (review criteria provided and no conflicting annotations; accessed August 2018). We turned to gnomAD^27^ to examine whether there are variants that are sufficiently common in the population to make it likely that they do not cause disease^15,22,27^. Of the 80 PTEN variants we analyse 29 after removing those absent from the experimental and computational data (Fig. 1). Of these, A79T and M205V are found at allele frequencies of 1·10^−4^ and 2·10^−5^, respectively, in gnomAD, that suggest they are likely benign.

We analysed the distribution of scores for the gnomAD and pathogenic variants as obtained by the four different methods (Fig. 2) and find that all four methods show a clear difference in averages and distribution of values across the gnomAD and pathogenic variants. We then performed a ‘receiver operating characteristic’ (ROC) curve analysis of each of the four methods two quantify their ability to separate pathogenic from the gnomAD dataset (Fig. 3). The ‘area under the curve’ (AUC) show that two experimental approaches perform better than the corresponding purely computational method (Fig. 3), and that the two methods that probe a wider range of functionally-relevant properties (Phosphatase-MAVE and evolutionary sequence energies) are able to capture variant effects more accurately than those that focus more directly on protein stability. Nevertheless, the high AUCs for the methods that probe only stability but not other aspects contributing to protein function (VAMP-seq and Rosetta) suggest that loss of stability and resulting decrease in cellular abundance is a key driver for PTEN-associated diseases in line with previous computational^4,29–31^ and experimental^18–21,32^ studies.

**Figure 2:**
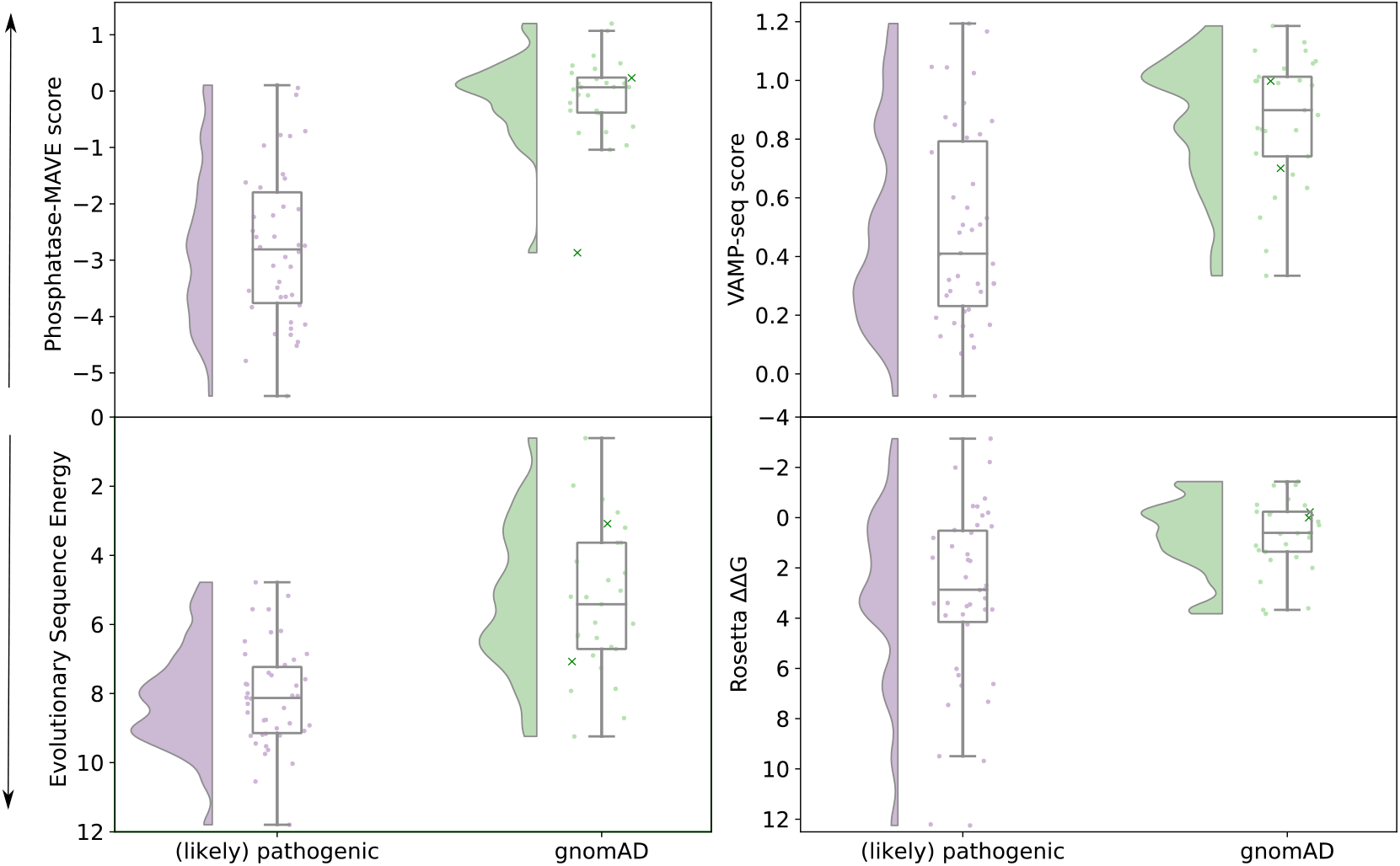
Distributions for pathogenic and gnomAD variants. Distributions of scores from each of the four methods for ClinVar pathogenic (mauve) and gnomAD (green) variants, shown as Raincloud plots^33^. Boxes illustrate the 25^th^-to-75^th^ percentile, with the median indicated by a horizontal line. Whiskers extend to minimum and maximum value though at most 1.5 times the distance between the 25th and 75^th^ percentile (“interquartile range”) away from the box boundaries. As described in the main text, pathogenic variants were removed from the gnomAD set. A79T and M205V are specifically highlighted (crosses), as these are the most common PTEN variants in gnomAD, and thus the most likely to be benign. Only variants for which each of the four methods provide data are included (see also Figure 5.1g). Axes are oriented such that values near the bottom correspond to detrimental variants, while values near the top are wild-type-like.

**Figure 3:**
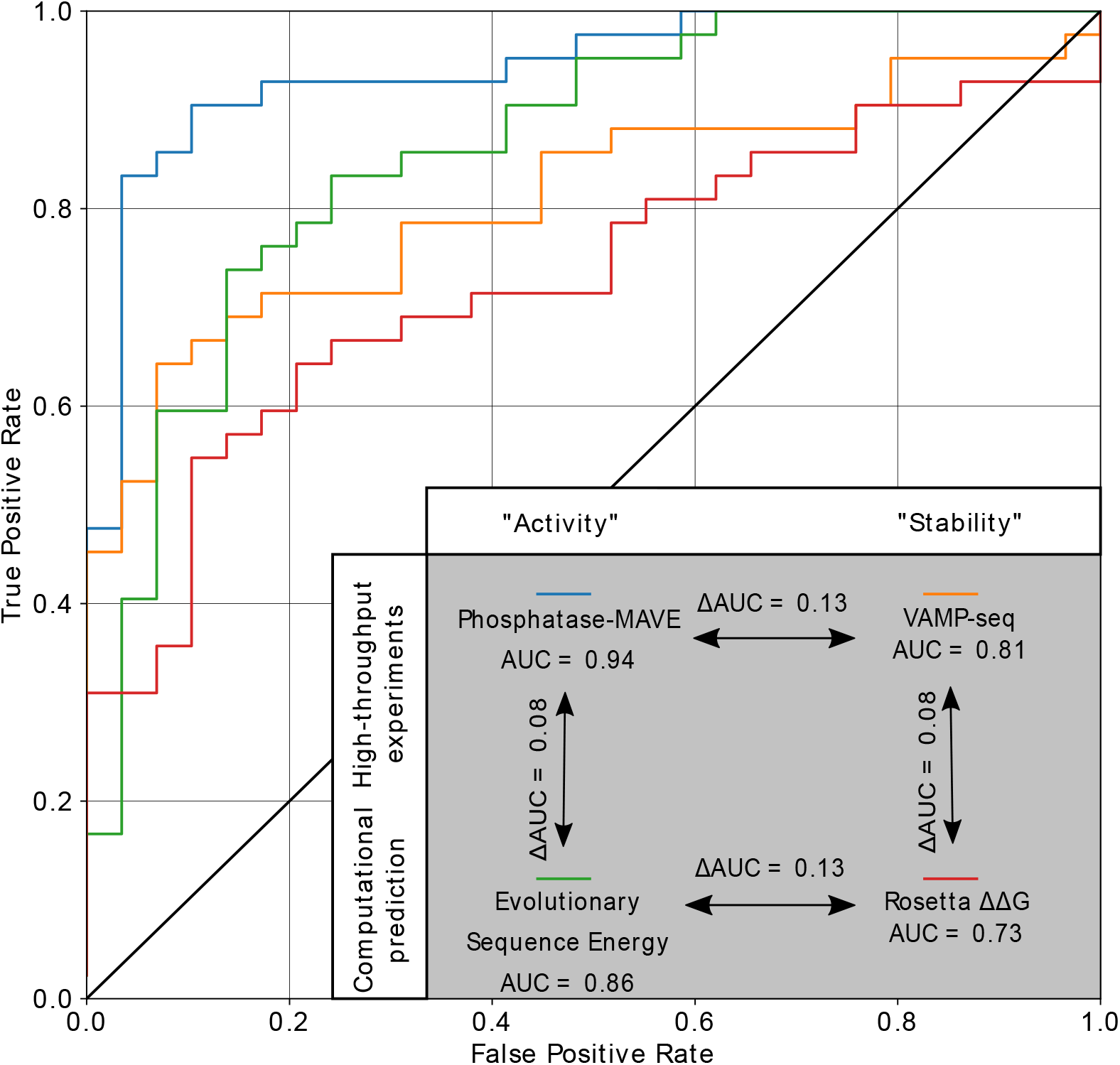
Receiver Operating Characteristic (ROC) curves for each method. ROC curves describe how well each method separates the ClinVar pathogenic variants from the gnomAD variants. As expected, methods assessing overall PTEN function perform better, with Phosphatase-MAVE experimental data (blue, AUC 0.94) exceeding evolutionary sequence energies (green, AUC 0.86) in overall sensitivity and specificity. Assessment of stability alone correctly identifies most pathogenic and gnomAD variants as well, again with experimental data (VAMP-seq, orange, AUC 0.81) exceeding performance of Rosetta ΔΔG calculations (red, AUC 0.73). The insert shows the AUC values and changes in AUC when moving between experimental and computational methods, and between methods that capture activity broadly or methods focused on stability.

We show two-dimensional plots of the change in stability (VAMP-seq or Rosetta (ΔΔG) and “activity effects” (Phosphatase-MAVE or evolutionary sequence energies) (Fig. 4) and separate the plots into four quadrants^19,21^ (Methods). Through this analysis we find that 27 of the 42 pathogenic variants fall in the unstable-and-non-functional quadrant in the experimental analysis (Fig. 4A) and 24 of the 42 variants fall in the corresponding quadrant in the computational analysis (Fig. 4B), suggesting that about 60% of disease-causing variants in PTEN cause disease via loss of stability and cellular abundance. The observation that most unstable variants are non-functional, but that not all non-functional variants are unstable is also reflected in the distribution of scores (Fig. 2) and ROC analysis (Fig. 3). Thus, if we examine the ROC curves in the region of high specificity (10% false positive rate) we reach a relatively high sensitivity (true-positive rate) for all methods (Phosphatase-MAVE: 0.90, VAMP-seq: 0.66, Gremlin: 0.66, Rosetta 0.53). Thus, one can find ~60% of the pathogenic variants at a relatively low false discovery rate, simply by examining loss of stability.

**Figure 4:**
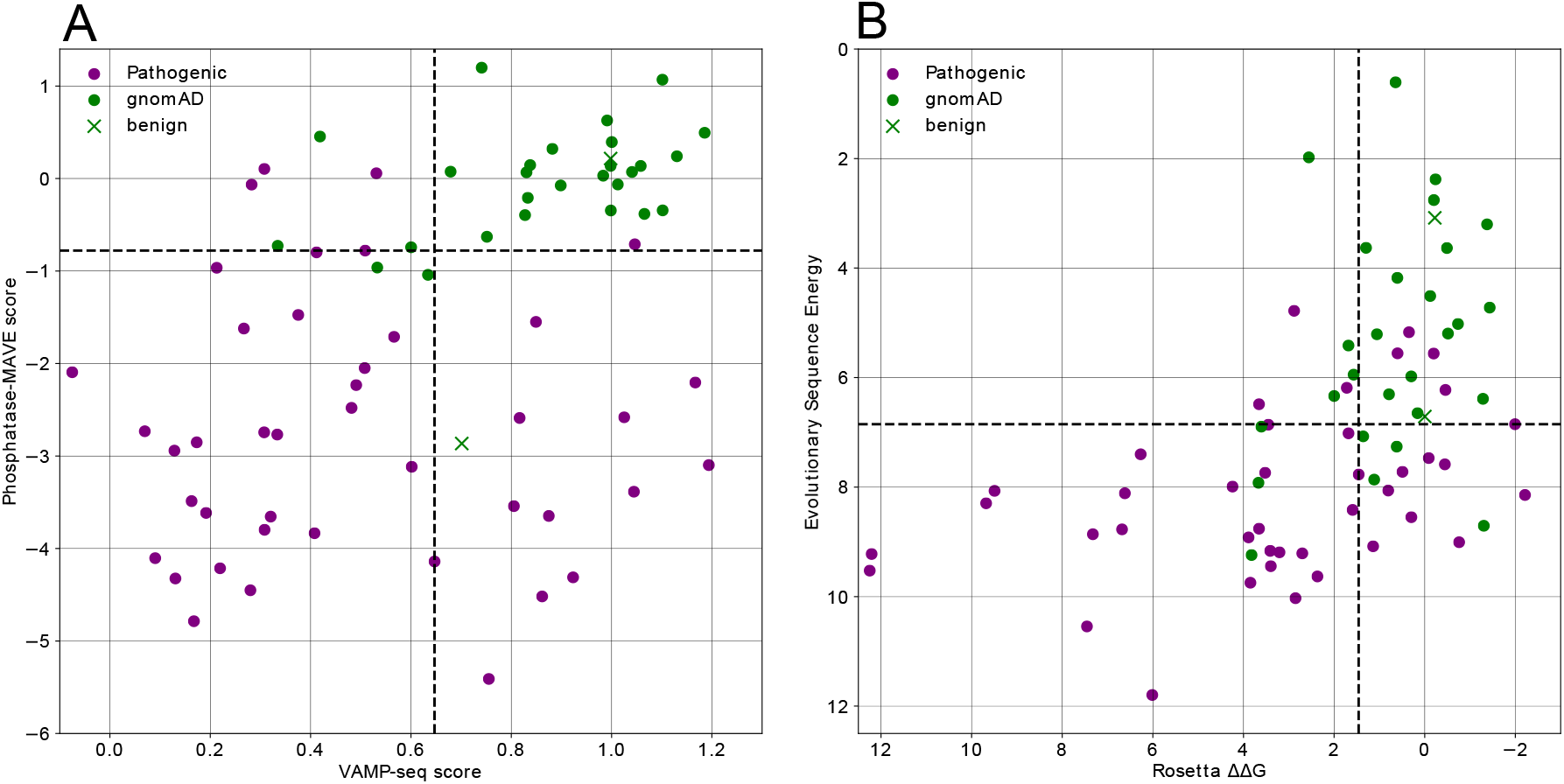
Two-dimensional landscapes integrating changes in stability and fitness. Combining scores from assessment of stability and function shows that most gnomAD variants (green) are wild-type-like, while most pathogenic variants (purple) are correctly identified by one, and often both metrics. Comparing the landscape based on experimental data (left) to that based on predictions (right) shows that availability of experimental data leads to better separation of pathogenic variants, but also that the predictions will provide a good starting point at substantially lower cost. The dashed lines correspond to cut-off values derived from the ROC curves, as those values that give rise to the point on the ROC curve closest to the upper-left corner (TPR=1, FPR=0).

## Conclusions

We have here taken advantage of the availability of both Phosphatase-MAVE and VAMP-seq data for the human protein PTEN to analyse how well these methods are able to classify known pathogenic variants, and have compared the results to computational methods. Our results show that all four methods we tested are able to provide relatively accurate classifications of variant effects with AUCs ranging from 0.74-0.94 (Fig. 3). Here we remind the reader that we in our selection of pathogenic variants have combined disease-causing variants for different diseases (ASD and PHTS) although it has been suggested that ASD results from variants with milder effects compared to those that give rise to PHTS^15^. Similarly, we note that we have used also rare variants from gnomAD as a proxy for benign variants, and that some of these may indeed give rise to ASD or PHTS. With these caveats in mind we highlight two observations regarding prediction accuracies. First, the experimental methods give rise to AUCs that are about 0.1 unit higher than the corresponding computational method (Phosphatase-MAVE (0.94) vs. evolutionary sequence energies (0.86) and VAMP-seq (0.81) vs. Rosetta (0.73)). Second, the methods that probe function more generally give rise to AUCs that are about 0.1 unit higher compared to those that probe only protein stability (Phosphatase-MAVE (0.94) vs. VAMP-seq (0.81) and evolutionary sequence energies (0.86) vs. Rosetta (0.73)). One advantage of the methods that focus on protein stability is that they provide a more direct mechanistic model for how the variants cause disease. Indeed, our two-dimensional analysis of stability and ‘function’ reveal that many, but not all, disease-causing variants appear to cause loss of function because the variants cause loss of stability and decreased cellular abundance (Fig. 4). We conclude that at the current stage, experimental methods still perform better than the corresponding computational approaches, although the latter perform well on many variants too. Further, we note that many, but not all, pathogenic variants in PTEN appear to give rise to disease via loss of stability and cellular abundance.

## Methods

### Rosetta ΔΔG Calculations

We used the “cartesian_ddg” application in Rosetta with the ‘beta_nov_16’ variant of the Rosetta energy function to perform the ΔΔG calculations^26^ using (PDB ID 1D5R) and Babel^34^ to parameterise the ligand. To accomodate for the missing loop in the crystal structure (residues 281 to 313), we added the flag “-missing_density_to_jump” during relaxation of the structure.

When calculating ΔΔG we used three iterations, which were subsequently averaged. The resulting difference in stability was finally divided by 2.9 to bring the ΔΔG values from Rosetta energy units onto a scale corresponding to kcal/mol (Frank DiMaio, University of Washington; personal correspondence).

### Evolutionary sequence energies 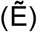

We used HHblits^35^ to create a multiple sequence alignment based on the UniProt sequence of PTEN (UniProt AC P60484) and used Gremlin^25^ to to calculate the log odds score between each single residue variant and the wild type.

### Phosphatase-MAVE and VAMP-seq Data

The VAMP-seq data can be accessed and downloaded at: https://abundance.gs.washington.edu/shiny/stability/.

The Phosphatase-MAVE data can be accessed in the supplementary material of the online article from Mighell et al.^15^

### ROC-derived thresholds

Thresholds for separating likely pathogenic from likely benign variants are calculated for each metric by determining the point on the ROC curve that is closest to (0,1), the optimum where perfect specificity and sensitivity would be achieved. In this case we give comparable weight to detection of pathogenic and benign variants, but note here that, depending on the application, different choices of threshold may be most suitable.

### Analysis scripts

The scripts used in the analyses and for making the figures of this manuscript are available at https://github.com/KULL-Centre/papers/tree/master/2019/PTEN-variants-Jepsen-et-al.

## Acknowledgements

We thank Dr. Sofie V. Nielsen, Mustapha C. Ahmed and Prof. Fritz Roth for helpful discussions and comments. This work is supported by a Novo Nordisk Foundation Challenge Grant (PRISM) (D.M.F, R.H-P., A.S., K.L-L.), the Lundbeck Foundation (A.S.), and the National Institute of General Medical Sciences (1R01GM109110 and 1RM1HG010461 to D.M.F.). D.M.F. is a CIFAR Azrieli Global Scholar.

